# Cooperative regulation of adherens junction elongation through Epidermal Growth Factor Receptor (EGFR) activation

**DOI:** 10.1101/2021.05.25.445561

**Authors:** Chaoyu Fu, Aditya Arora, Wilfried Engl, Michael Sheetz, Virgile Viasnoff

## Abstract

The mechanisms controlling the elongation rates of adherens junctions are poorly characterized and lag behind our understanding of those controlling their static properties. In suspended cell aggregates, we found that the speed of *de novo* junction formation between two cells increases with the number of junctions that the cells are already engaged in. This cooperative effect is driven by the transient activation of the Epidermal Growth Factor Receptor (EGFR) upon junction formation. EGFR activation then regulates the actin cytoskeleton turnover while cortical tension remains unaltered. Overall, we show that EGFR activation regulates the elongation speed of junctions (kinetype) without affecting their final size (phenotype) in such aggregates.

## Introduction

In the last decade, the nature and organization of the molecular components constituting adherens junctions have been under intense investigation. Much less is understood about how cells control the dynamics of adherens junction formation and remodeling. Cadherins, and in particular E-cadherin are capable of orchestrating the assembly of proteinaceous clusters involving actin-binding protein (eg: catenins), scaffolding proteins (eg: α-actinin,cortactin), and tight junction proteins (eg: ZO proteins) (Helwani et al., 2004; Huveneers and de Rooij, 2013). E-cadherin adhesion is also recognized as a signalling hub where RTK receptors (EGFR) and other GTPases (Rac1 and Cdc42) are activated (Fedor-Chaiken et al., 2003; Kovacs et al., 2002; Nakagawa et al., 2001; Noren et al., 2001; Pece and Gutkind, 2000). We have a fair understanding of how junctional components organize the mesoscale adhesive plaque and respond to mechanical stimulations. The mechanical forces and in particular tension emerged as an essential parameter to stabilize junctions. It results that myosin II and its regulatory pathways are critically involved to control junction size and stability (Priya et al., 2015; Ratheesh et al., 2012). In epithelia where adherens junctions have been analyzed, junction morphologies are governed by the local equilibrium of mechanical tension at the junction vertices as demonstrated by laser ablation (Rauzi et al., 2008).

Our understanding of junction remodeling arises principally from the studies of the paradigmatic *Drosophila* embryo epithelia where again actomyosin contractility plays a critical role to shrink or expand the junction (Bertet et al., 2004; Rauzi et al., 2008). Pulsatile apical contractions (Jessica and Fernandez-Gonzalez, 2016) and localized E-cadherin cluster recruitment (Rauzi et al., 2010) have, for example, been proposed to drive cell-cell contact exchange, but they are often characterized with respect to the final junction size and tissue organization. This tension-centric view focuses on the driving force of the junction elongation. However, the junction elongation speed is very likely governed by the intrinsic dynamics of the actin cytoskeleton reorganisation, potentially equivalent to the modulation of the junction viscosity (Clement et al., 2017).

This study proposes that in the context of suspended cells, the direct or indirect activation of EGFR during adherens junction formation regulates the actin turnover rate of the free cortex and indirectly the elongation speed of another junction involving the same cell. We used minimal systems of cell doublets, triplets, and quadruplets to precisely image the dynamic of elongations of the junction in stereotypical conditions. We found that junction elongation rates depended on the number of adherens junctions that the two adhering cells have already engaged prior to new contact formation. This resulted in a cooperative effect that accelerated the formation of multi-junctions. We then demonstrated that the actin turnover of the cell-free cortex prior to the junction formation was the important factor that controlled the elongation rate of the adherens junction. We determined that the activation of EGFR signalling by E-cadherin during junction formation resulted in a decrease of cortical actin dynamics and thus controlled the cooperative effect of junction formation.

## Results

### Adherens junction elongation rate scales with the number of pre-existing cell-cell contacts

In order to monitor the kinetics of the formation of adherens junctions, we used suspended cell doublets. The approach offers the advantages of *i*-precise timing the onset of junction formation, *ii*-creating round junctions with highly stereotypical quantifiable morphologies, *iii*-precluding any crosstalk from extracellular signaling by matrix. In a previous study (Engl et al., 2014), we established that this approach leads to a *bona fide* and stereotypical distribution of junctional proteins (E-cadherin, actin, vinculin, and catenins) along with the contact between two cells. Here we adapted the approach to analyze the kinetics of junction formation of S180-E-cad-GFP cells, a murine sarcoma cell line expressing no cadherins endogenously but stably transfected with E-cad-GFP (Chu et al., 2004). The results obtained on S180 were qualitatively reproduced using MDCK cells (see Supplementary Information). Nonetheless, MDCK cells are more difficult to handle as suspended aggregates due to rapid anoikis when manipulated in suspension.

We seeded suspended S180 cells inside non-adhesive microwells for fast live *en face* imaging of junction formation. Using custom Matlab image analysis, we measured the junction radial growth (Fig.1a and Supplementary Fig.1a, b). As previously described, GFP-E-cad accumulated at the junction rim into distinctly spaced clusters during the latter part of the junction formation process (Supplementary Fig. 1b) (Engl et al., 2014). Once an initial adhesion zone was established between two cells, the evolution of the radius R(t) of the circular contact served as an unambiguous proxy to measure junction dynamics (Figure 1a, b) since the elongations followed a well defined exponential relaxation dynamics towards the final radii.

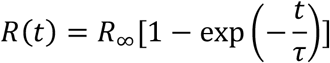

**Figure 1.**
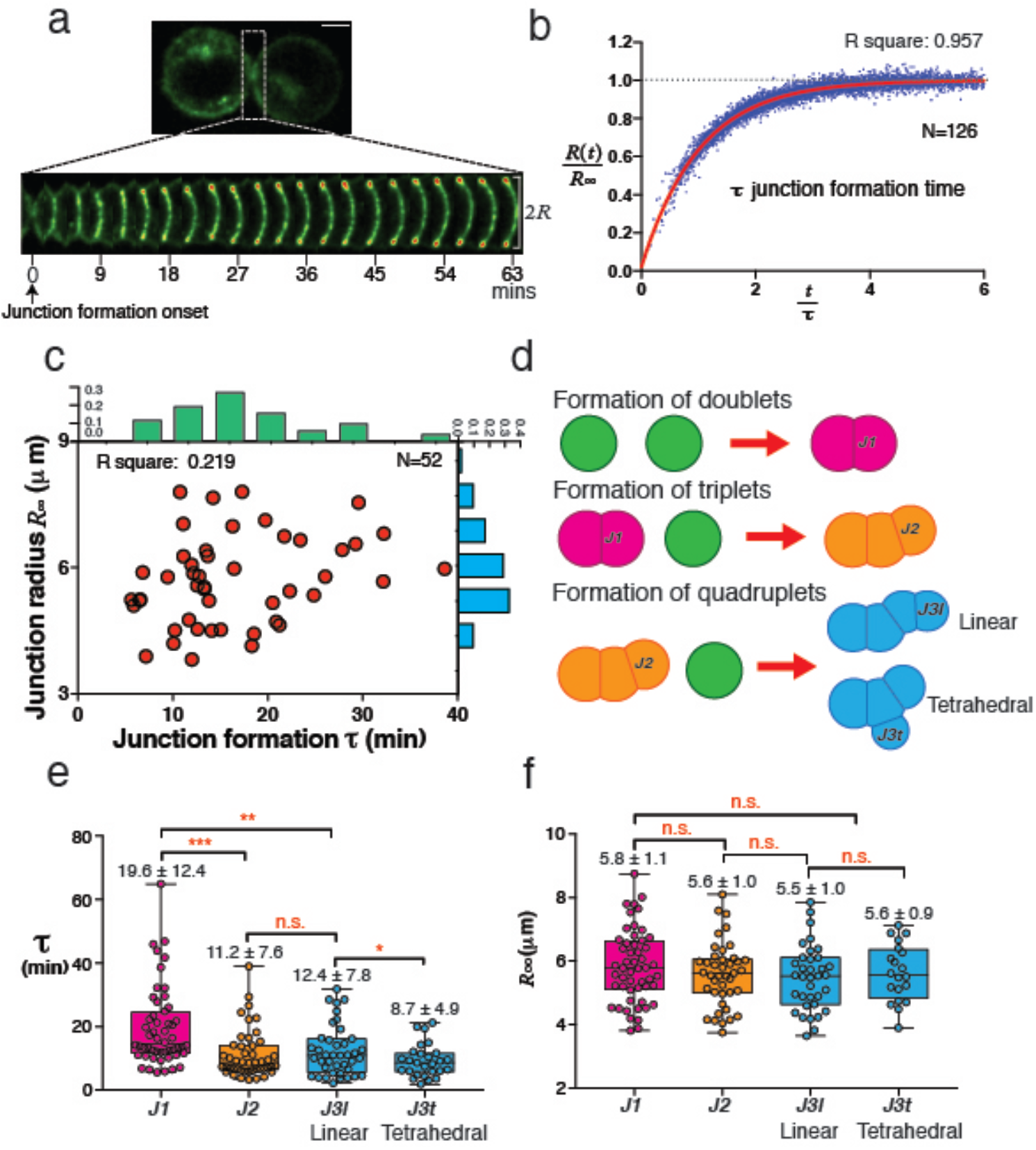
*De novo* adherens junction formation primes cells for the faster formation of subsequent adherens junctions. **(a)** Side view of the process of *de novo* adherens junction formation after junction initiation. Time interval, 3 mins. Scale bar, 5μm. **(b)** The evolution of junction radius in the function of time. Junction formation time (τ) is used to describe the junction formation speed. N=126. **(c)** Scatter plot of junction formation time and junction radius (red); Histogram of junction formation time (green) and junction radius (blue) for *de novo* adherens junctions. N=52. **(d)** Schematic side view of the formation of doublets (J1), formation of triplets (J2) and formation of quadruplets (J3) (Linear and Tetrahedral). **(e and f)** Box plots of junction formation time (τ) **(e)** and junction radius **(f)** in doublets (J1), triplets (J2) and quadruplets (J3) (Linear and Tetrahedral). n = 52 doublets, 46 triplets, 40 quadruplets (Linear) and 25 quadruplets (Tetrahedral) from three independent experiments. The P values in **e** and **f** are calculated from one-way ANOVA.

Surprisingly, the fit remained excellent under the various drug treatments used in this study as shown by Figure 1b where 126 cells and 6 conditions can be rescaled by their static sizes and dynamics onto a master exponential relaxation (R^2^=0.957). We concluded that the measurement of *R*_∞_ and *τ* sufficed to fully characterize the junction formation dynamics.

In control conditions, the elongation times of doublet junctions (J_1_) proved uncorrelated with the final junction size (Fig. 1c). Using calibrated confocal imaging we investigated the influence of E-cadherin density on the junction formation. We found that junctions formed provided that at least one cell of the doublet had a density higher than a threshold value (Supplementary Fig. 1c, d). The formation speed and the size of the contact, however, did not correlate with the level of cadherin expression (Supplementary Fig. 1e). The characteristic formation time of a junction in the control case was 19.6±12.4 min with a final radius of 5.8±1.1 μm. These observations suggested that the dynamic properties of a junction might be decoupled from its static properties.

We then asked whether a second junction (J_2_) with either cell of a doublet would elongate at the same rate. Contacting doublets with singlets (see Materials and Methods), we hence formed linear triplets (Fig. 1d and Supplementary Fig. 1f). We followed the junction formation dynamics of triplets that were seeded in the same microwell. In linear triplets, the second adherens junction (J_2_) (11.2±7.6 min) formed 37 % faster than the first adherens junction (J_1_) (Fig. 1d, e and Supplementary Fig.1f-h). It is important to note that we considered exclusively the dynamics of elongation once initial contact is formed hence excluding the delay needed for cells to meet and to initiate the contacts. The observed increase in the elongation rate for J_2_ did not depend on the time elapsed between the initiations of J_1_ and the J_2_ (within 1hour) (Supplementary Fig. 1i, j). Older doublets did not show any speed up of junction elongation when engaged in triplets, hence suggesting a transient priming by the formation of J_1_. We observed a similar effect with MDCK cell doublets and triplets, indicating this effect was not S180 specific (Supplementary Fig. 1k).

We then analyzed the formation of a third junction to form cell quadruplets. For linear quadruplets when the third junction (J_3l_) formed with either end cell of the triplet, J_3l_ elongated at the same speed as J_2_ (12.4±7.8 min) (Figure 1e). By contrast, when the third junction (J_3t_) formed with the central cell of the triplet, J_3l_ elongated 22% faster than J_2_ (8.7±4.9 min) and consequently 56% faster than J_1_ (Figure 1e). In the tetrahedral morphology, the third junction involves the central cell that is already engaged in two contacts whereas in linear quadruplets the end cells are engaged a single pre-existing contact. It consistently indicates that the elongation speed of a junction, in this context, increases with the number of pre-existing junctions. Remarkably, the final radii of the junctions J_1,_ J_2,_ J_3l,_ J_3t_ were equivalent (Fig. 1f). Taken together our data indicate that the dynamics of junction formation is cooperative but largely decoupled from the final junction morphology.

### EGFR activation after *de-novo* junction formation regulates the cooperative dynamics of junction formation

In the absence of extracellular matrix adhesion, we reasoned that the cooperative effect was triggered solely by the *de novo* junction formation. In particular, we hypothesized that the engagement of E-cadherin during the first junction formation triggered a change of the actin cytoskeleton constituting the free cortex of the cell. We reasoned that the parameter driving the junction elongation was the dynamics of actin restructuring from the free cortex into the junctional actin. We considered mechanical tension and actin turnover rate as the two main potential cortex biophysical properties that the cells could regulate to alter contact elongation. We hence monitored the tension of the free cortex by pipette aspiration and its actin turnover rate using Fluorescence Recovery after Photobleaching (FRAP) as detailed in Materials and Methods (summarized in Figure 2A). We compared junction and cortex properties for singlets, doublets, and triplets. First, we found that single cells or cells engaged in contact had the same free cortical tension at early times (<10 minutes) and at long times (>1h) alike (Fig. 2b, and Supplementary Fig. 2a). This suggested that the formation of the junction did not enhance the contractility (Myosin activity) of the cell cortex. In contrast, the average actin recovery time at the free cortex (t_Cort._) displayed an 87% increase (from 10.9±4.3s to 20.4±7.3s) between a population of singlets and doublets (Fig. 2c, Supplementary Fig. 2b-d). This holds true at the single-cell level, as probed by repeated FRAP measurements of individual cortices before and after *de novo* junction formation. At the single-cell level, the turnover recovery time t_Cort_ systematically increased, hence matching the conclusions from the population study. (Supplementary Fig. 2e). It suggested that the E-cadherin junction formation could trigger a signaling pathway that rapidly altered the dynamical properties of the free cortex cytoskeleton. We concluded that cell multiplets can have a similar static phenotype (Number of contacts, size of contact, cortical tension) but distinct kinetypes (actin turnover dynamics, junction formation speed). It suggested that some distinct pathways could be involved to independently regulate the dynamical properties and the static properties of adherens junctions.

**Figure 2.**
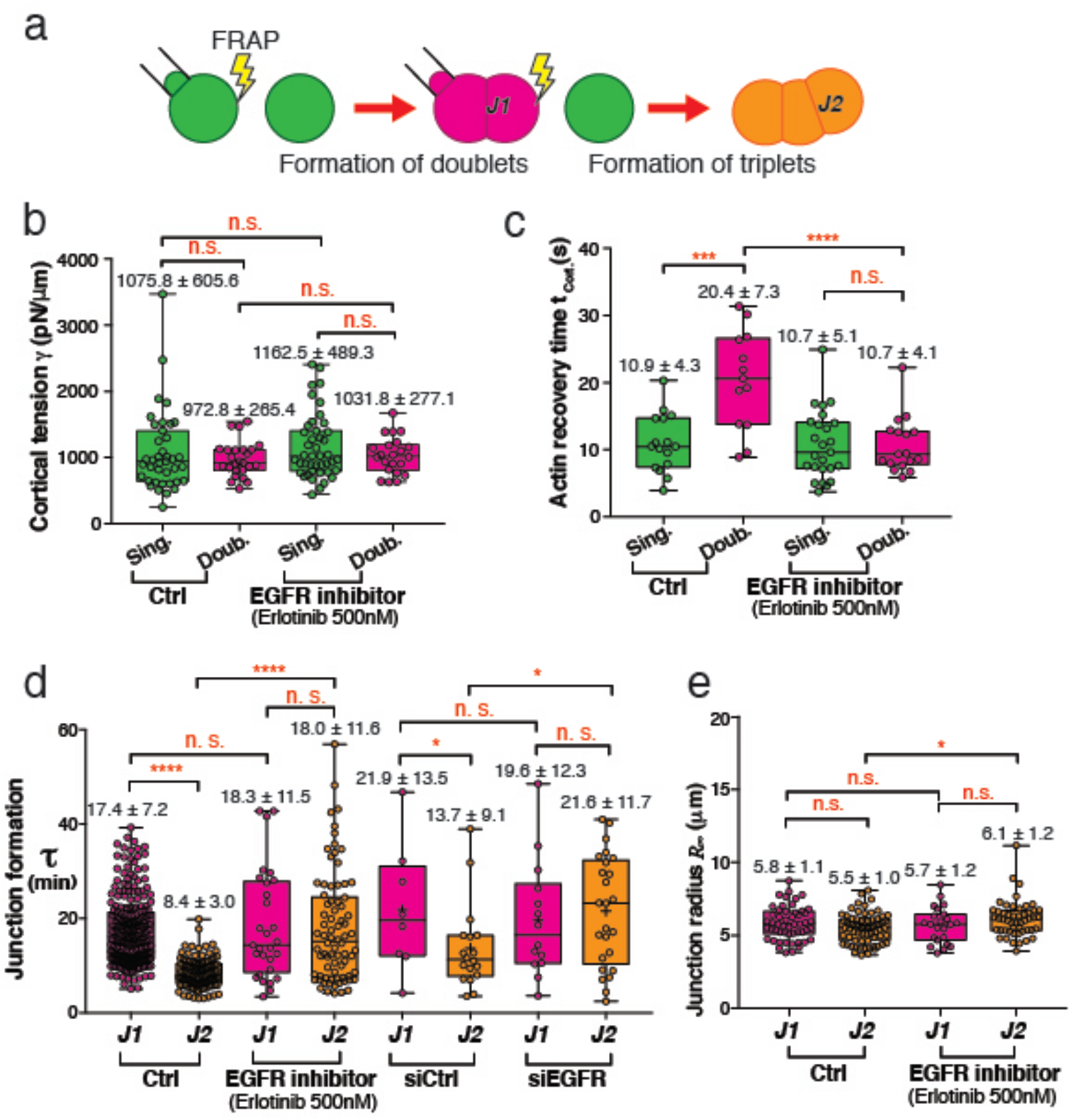
The cooperative effect is regulated by EGFR activity through altering cortical actin dynamics after *de novo* junction formation. **(a)** Schematic of FRAP and micropipette aspiration measurements in the cortex of singlets and doublets and the side view of the formation of doublets (J1) and formation of triplets (J2). **(b and c)** Cortical tension **(b)** and actin recovery time **(c)** for singlets cortex and doublets cortex. (**(b)** Control, n= 16 singlets and 14 doublets, treated with Erlotinib at 500nM, n= 25 singlets and 17 doublets; **(c)** Control, n= 39 singlets and 25 doublets, treated with Erlotinib at 500nM, n= 43 singlets and 22 doublets). Junction formation time for doublets (J1) and triplets (J2) (Control, n= 238 doublets and 115 triplets, treated with Erlotinib at 500nM, n= 29 doublets and 78 triplets, nontarget siRNA control, n= 8 doublets and 18 triplets, siRNA against EGFR, n = 14 doublets and 25 triplets). **(e)** Junction radius for doublets (J1) and triplets (J2) (Control, n= 52 doublets and 72 triplets, treated with Erlotinib at 500nM, n= 23 doublets and 50 triplets). The *P-*value in **b, c, d and e** are calculated from one-way ANOVA.

Hence, we searched for pathways that can alter the kinetype of the cell junction dynamics while preserving the phenotype. E-cad engagement can reportedly lead to the phosphorylation of several kinases (McLachlan et al., 2007; Pece and Gutkind, 2000). Using phospho-specific antibodies, we screened several Receptor tyrosine kinases (RTKs). We found that EGFR was phosphorylated at the Src-dependent domain Y845 at the junction edge (Supplementary Fig. 3). We then analyzed the cortical actin dynamics in singlets and doublets after pharmacological inhibition of EGFR (Erlotinib at 500nM, see Materials and Methods for specificity). Inhibition of EGFR activity *i*-did not alter the cortical tension of singlets and doublets (Fig. 2b), *ii*-fully inhibited the slowdown of the actin turnover dynamics t_Cort._ upon junction formation (Fig. 2c and Supplementary Fig. 2e), *iii*-blocked the cooperativity of junction formation between J_1_ and J_2_ (also confirmed by siRNA partial KD) (Fig. 2d and Supplementary Fig. 2f), *iv-* did not impact the final radius of the junctions (Fig. 2e and Supplementary Fig. 2g). Identical conclusions applied when the inhibitor was used during cell quadruplets (J_3l_ and J_3t_) formation (Supplementary Fig. 2h, i). As a whole EGFR inhibition did not alter the junction phenotypes in the cellular multiplets but abolished the difference in their kinetypes and hence the cooperative effect of junction elongation.

These correlative observations prompted us to postulate that the alteration of actin turnover downstream of EGFR activation upon E-cad junction formation could result in accelerating the formation of subsequent junctions.

### Slower turnover of the free actin cortex speeds up junction elongation

We then examined if a modification of cortical actin turnover could cause a change in junction elongation rate independently of its molecular origin. We used a series of pharmacological inhibitors (latrunculin (250nM), CK-666 (100μM), SMIFH2 (20μM), and jasplakinolide (100nM)) that target different mechanism of actin regulation (polymerization, branching, depolymerization). We performed systematic FRAP and cortical tension measurements on singlets and correlated them with the speed of elongation of the doublet junction (J_1_). We compared all kinetypes and phenotypes under the various treatment conditions. Supplementary Figure 4 displays all the six combinations of correlative graphs relating the possible pairs of parameters cortical tension γ, cortical actin recovery time t_Cort._, final junction radius *R*_∞_, and junction formation time *τ*.

The actin dynamics of the free cortex t_Cort._ proved largely decorrelated from *R*_∞_ and *γ* (Supplementary Fig. 4a, c). The junction formation time τ displayed its strongest correlation (R^2^=0.655) with t_Cort._, irrespective of the drug treatment (Supplementary Fig. 2b). It also displayed a weaker correlation with the cortical tension *γ* (Supplementary Fig. 2d). These results concurred with what we described for doublets, triplets, and quadruplets. They confirmed that the modulation of the actin dynamics alone can cause, directly (mechanosensitive effect) and/or indirectly (reduction of protrusion rate, altered signalling), the change in the speed of elongation of a junction. They reinforced our hypothesis that the alteration of actin turnover at the free cortex upon EGFR activation can causally change the junction dynamics.

We further explored how EGFR activation could regulate actin dynamics. The cooperative formation of junctions proved insensitive to the presence of EGF (150nM) in the cell culture medium (Fig. 3). Indeed, the effect was quantitatively similar between control and serum-starved cells (+serum: 17.4±7.2 min and 8.4±3.0 min, -serum: 19.6±12.4 min and 11.0±7.0 min) (see Materials and Methods for the detailed protocol). In both cases, the inhibition of EGFR abolished the systematic increase of junction elongation speed between J_1_ and J_2_ (19.0±11.1 min and 16.3±11.1 min) (Fig. 3a, b) as well as the alteration of actin dynamics (11.1±5.6 s and 11.9±4.7 s) (Fig. 3a, c). It suggested that the effect was not dependent on the basal level of activation of EGFR but could instead result from a transient burst of activation upon contact formation. To test if a pulsed activation of EGFR could alter the junction formation dynamics, we added EGF at 150nM to serum-starved singlets or doublets. As mentioned before, prolonged exposure to EGF (>1h) did not affect the cooperative effect as compared to control. However, when EGF was swiftly added just prior to the junction formation (< 10min), the elongation dynamics of the first junction J_1_ increased as compared to control (9.2 ±7.2 min, vs 19.6±12.4 min) to reach the same level as J_2_ in presence or absence of EGF burst (10.6 ±5.5 min and 8.4 ±3.0 min, respectively) (Fig. 3b). The actin recovery time in all three conditions also levelled to the slow t_Cort._ (Fig. 3c). We concluded that the burst of EGFR activation can induce a kinetype of the first junction that matches that of the second junction. As a whole, our data indicate that EGFR inhibition abolished the cooperativity effect of junction elongation levelling all junction to the slow kinetype of J_1_. By contrast, transient burst activation of EFGR by EGF also abolished the cooperative effect levelling all junction kinetypes to the fast one (J3). Our data hence strongly a transient activation of EGFR by junction formation that leads to a change in actin dynamics in the free cortex and consequently an increase in elongation speed. The activation could be ligand-independent as previously reported upon E-cadherin engagement (Fedor-Chaiken et al., 2003; Pece and Gutkind, 2000; Shen and Kramer, 2004) or mediated by a fast paracrine secretion of EGF by the cell.

**Figure 3.**
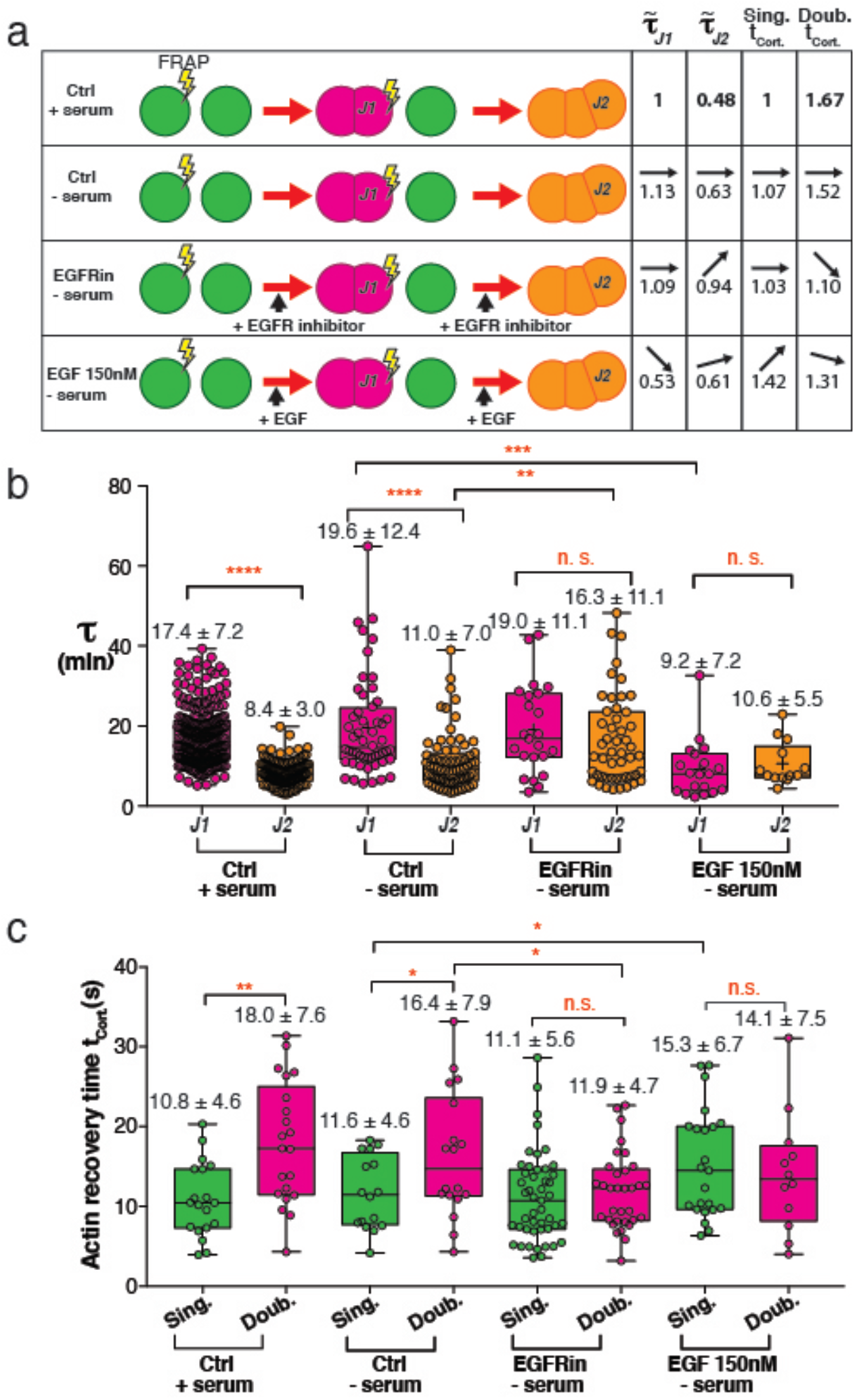
Burst of EGFR activity by adherens junction formaton determines the cooperative effect. **(a)** Schematic of junction formation of doublets (J1) and triplets (J2) in normal condition, serum-starved condition, serum-starved cells in the presence of EGFR inhibitor and EGF. Junction formation time and actin recovery time are normalized for comparison. **(b)** Junction formation time for doublets (J1) and triplets (J2) in the conditions listed in **(a)**. (control condition, n= 238 doublets and 115 triplets, control with serum starvation, n= 52 doublets and 72 triplets, treated with Erlotinib at 500nM, n= 23 doublets and 50 triplets, treated with EGF at 150nM, n= 19 doublets and 13 triplets). **(c)** Actin recovery time for singlets cortex and doublets cortex. (control condition, n= 16 singlets and 13 doublets, control with serum starvation, n= 16 singlets and 18 doublets, treated with Erlotinib at 500nM, n= 47 singlets and 33 doublets, treated with EGF at 150nM, n= 23 singlets and 12 doublets). The *P-* value in **b** and **c** are calculated from one-way ANOVA.

### The cooperative effects in junction elongation involves Src activity and Rac-Arp2/3 pathway

The EGFR phosphorylation site (Y845), which we found activated at adherens junction, is a c-Src-dependent activation site. Both c-Src and Rac were reported to be activated by E-cad adhesion during junction formation, where the Rac activity required the activation of EGFR (Betson et al., 2002; McLachlan et al., 2007). We hence studied the role of Src and Rac activity in regulating the cooperative effects. We found that using pharmacological inhibitors to inhibit Src and Rac1 activity both blocked the cooperative effect and the slowdown of cortical actin dynamics after junction formation (Fig. 4). Upon inhibition of Src and Rac1, junction formation time of doublets remained unchanged but the junction formation time of triplets was increased by 115% and 105%, respectively (Fig. 4a). Actin recovery time (t_Cort._) was reduced by 22% and 34% in doublets while having no change on singlets (Fig. 4b). According to previous reports that c-Src was required for E-cad adhesive interaction in CHO cells (McLachlan et al., 2007), we also found that Src kinase activity was involved in regulating junction size (Supplementary Fig. 5a). We further elaborated the pathway by focusing on Arp2/3, a well-known effector downstream of Rac1. Consistent with previous results, inhibiting Arp2/3 reduced the actin recovery time (t_Cort._) by 18% in doublets while having no effect on singlets (Fig. 4b). Arp2/3 inhibition also abolished the systematic increase of junction elongation speed between J_1_ and J_2_ (Fig.4a). These results suggested that the slowdown of cortical actin dynamics after *de novo* junction formation and the cooperative effect likely depended on Src activity and Rac-Arp2/3 pathway.

**Figure 4.**
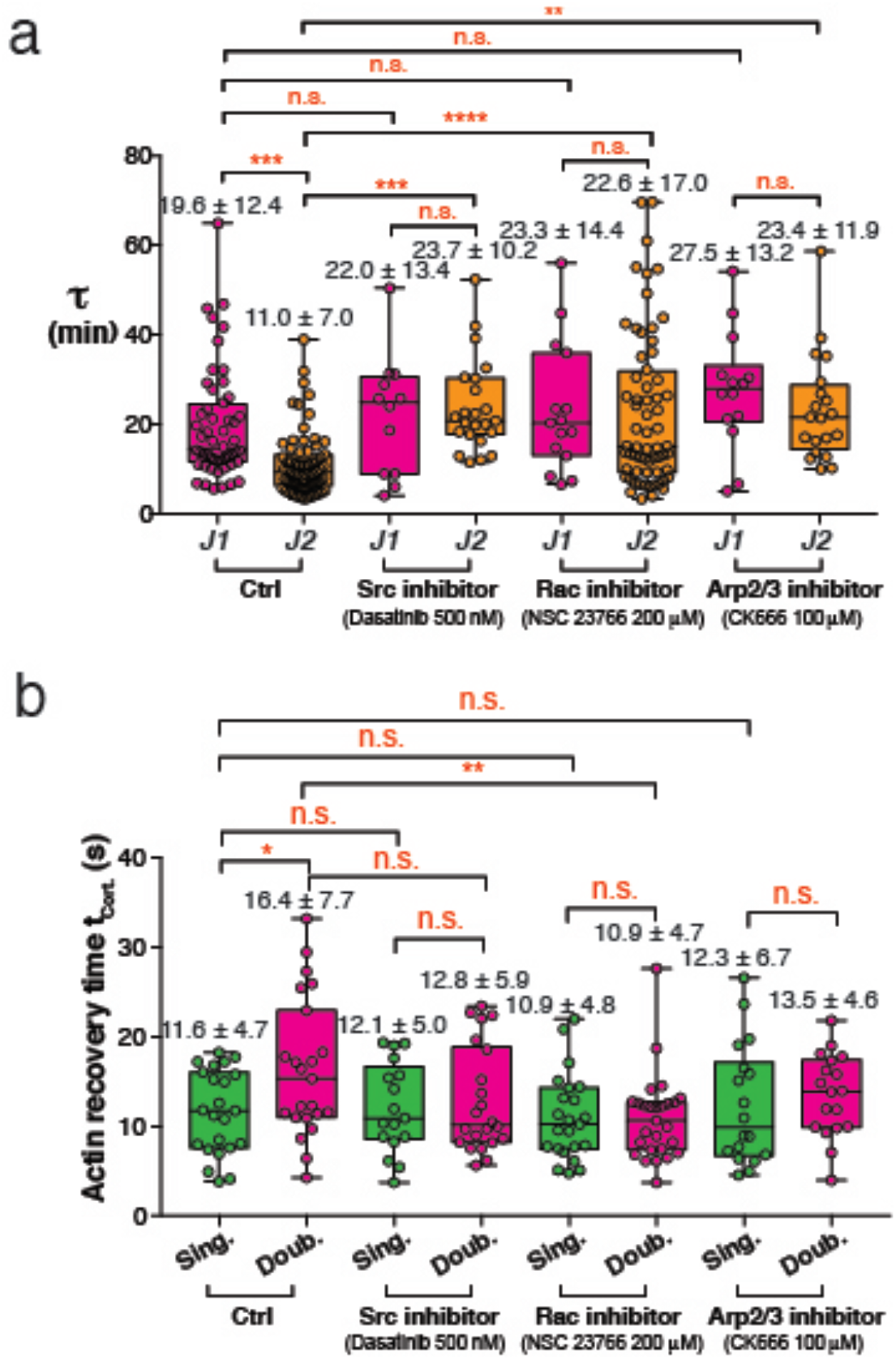
Slowdown of cortical actin dynamics after *de novo* junction formation and the cooperative effect depend on Src activity and Rac-Arp2/3 pathway. **(a)** Junction formation time for doublets (J1) and triplets (J2) in the serum starved conditions, treated with Src inhibitior, Rac inhibitor and Arp2/3 inhibitor. (control with serum starvation, n= 52 doublets and 72 triplets, treated with Dasatinib at 500nM, n= 12 doublets and 23 triplets, treated with NSC23766 at 200μM, n= 15 doublets and 64 triplets and treated with CK666 at 100μM, n= 14 doublets and 20 triplets). **(b)** Actin recovery time for singlets cortex and doublets cortex. (control with serum starvation, n= 23 singlets and 23 doublets, treated with Dasatinib at 500nM, n= 17 singlets and 22 doublets, treated with NSC23766 at 200μM, n= 23 singlets and 27 doublets, treated with CK666 at 100μM, n= 18 singlets and 18 doublets). The *P-*value in **a** and **b** are calculated from one-way ANOVA.

## Discussion

The existence of a cross-talk between adherens junctions and EGFR has been documented in many contexts. The general understanding is that prolonged EFGR activation by EGF destablisizes adherens junctions in epithelial cells, promotes E-cadherin endocytosis, and triggers mesenchymal transition of epithelial cells (Barrandon and Green, 1987; Lu et al., 2003). At the same time, polarized epithelium is less sensitive to EGF due to the sequestration of EGFR at the lateral pole beneath the zonula adherens, leading to reduced exposure to the soluble ligand (Kim et al., 2009; Perrais et al., 2007; Qian et al., 2004). This mechanism contributes to the resistance of epithelium to metastasis. In cancer, the soluble ectodomains of E-cadherin, cleaved from cancer cells into the surrounding medium are also thought to promote activation of EGFR and consequently destabilisation of the junction (Rodriguez et al., 2012). Recent works have highlighted and unraveled an interdependent relationship between RTKs (especially EGFR) and cellular cytoskeletal networks. For example, Roth et al. identified LAD-1 as a target of EGFR activation which plays a role in regulating actin dynamics that is critically involved in cell migration (Chiasson-MacKenzie and McClatchey, 2018; Roth et al., 2018). Further, magnetic twisting cytometry (MTC) was used to reveal that E-cadherin-based force transduction leads to cell stiffening through the activation of EGFR (Muhamed et al., 2016).

Our doublet system here establishes, at least for this cell model four main points that differ from previous literature reports: *i-* EGFR can be activated directly or indirectly by the *de novo* formation of E-cadherin junctions in absence of integrin signaling. *ii-* This fast and transient activation leads to a change in the properties of the whole cell cortex and not only the junction itself. *iii-* The free cortex in the activated EGFR cells displays slower actin turnover, probably originating from the activation of Arp2/3, which leads to a faster elongation. *iv-* A short burst of EGFR activation by soluble EGF (but not a prolonged exposure to EGF) can phenocopy the priming by *de novo* adherens junction formation (summarized in Figure 5). It suggests that the activated signaling pathway downstream of the phosphorylation of EGF might be selected by the kinetics of the activation. Moreover, our data strengthen the observations that Arp2/3 activity leads to a decrease in actin turnover while formin activation will increase actin dynamics. These results were in line with the previous report that the actin assembly speed is decreased in mDia1-depleted cells while perturbation of Arp2/3 leads to an increase (Bovellan et al., 2014). This phenomenon is caused by the competitive regulation of actin dynamics by formin and Arp2/3. The author found that actin assembly speed (dynamics) increases if a slower-than-average nucleator (such as Arp2/3) is depleted while depleting a faster-than-average nucleator (such as mDia1) will decrease actin assembly speed (dynamics) (Bovellan et al., 2014).

**Figure 5.**
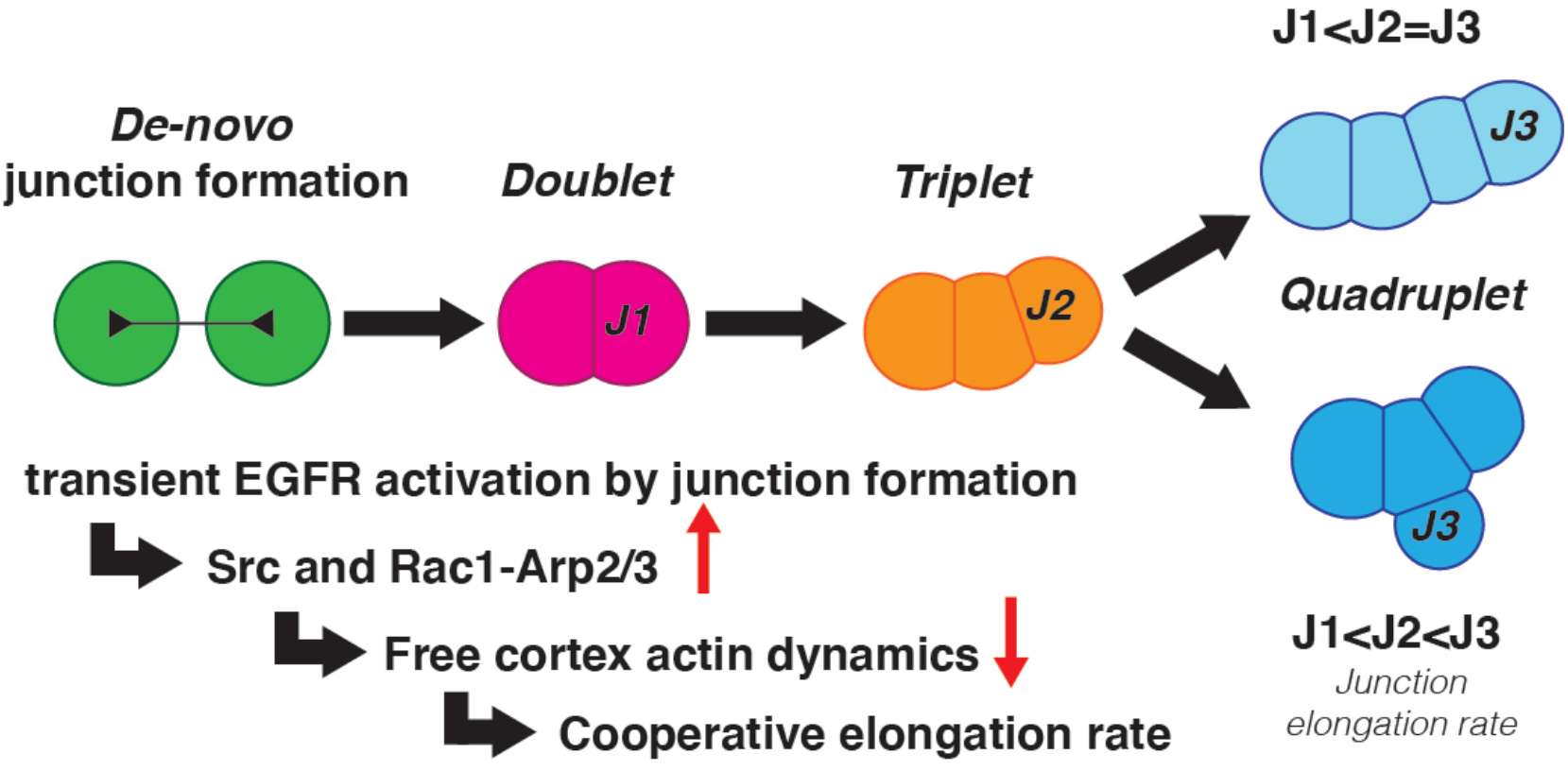
Proposed model for the cooperative effect among junction formation, EGFR activation and actin dynamics modulation that promotes rapid formation of multiple cell junctions. **(a)** Initial junction formation leads to EGFR activation. This EGFR activity slows down cortical actin dynamics distal from the junction. Low actin dynamics primes cells for faster junction formation to form multiple cell junctions.

To our knowledge, this study is first to establish that the junction elongation speed can be controlled by cells through an RTK activation that impinges on global cortex properties. It is important to note that the transient activation of EGFR alters the junction elongation speed but is not necessary to trigger junction formation. This could be either due to alternative or redundant regulation of the cortex by other RTK or regulatory pathways (Bedzhov et al., 2012). It could also be that EGFR activation is not involved in the early stage of contact formation.

Last but not least, we would like to emphasize that our study showed that the dynamical properties of the junctions in our systems (kinetypes) are decoupled from the final size of the contact (phenotype) that is governed by the level of contractility of the cortex.

At this stage of the study, we can only speculate as to why the junction formation dynamics depend on actin turnover more than on contractility. Our data suggest that, at least in the context of suspended cells, cortical tension determines the size of the contact, in line with the current understanding that junction size results from the mechanical balance of cortical and junctional tension. The simple assumption that the junction elongation results from cortical tension acting against a dissipative viscosity of the created interface and free cortex fails to account for the exponential relaxation that robustly describes all our data. At the moment we do not have a satisfying model to account for the central role of free cortex dynamics in the elongation rate of the junction. However we could hypothesize that the elongation rate is controlled either by *i*-the dynamics of rearrangement of actin at the edge of the junction, or by *ii*-the local protrusive activity of the free cortex in the close vicinity of the contact edge. Junction elongation in this context would echo, at least locally the zippering mechanism invoked in keratinocytes or in fibroblasts. Further theoretical studies are needed to precisely model the observations.

Our study raises the interesting point that cells can regulate independently the dynamics and the static outcome of their contact. Acto-Myosin contractility can thus not be considered as the sole parameter that controls the junction evolution.

## Materials and methods

### Antibodies and reagents

Antibodies and pharmacological inhibitors were as follows: rabbit anti-phospho-EGFR (Y845) polyclonal antibody (Invitrogen # 44-784G, 1:200 dilution); rabbit anti-EGFR antibody (Cell Signaling Technology #2232S); mouse beta Actin Loading Control Monoclonal Antibody (BA3R) (Invitrogen #MA5-15739); anti-Rabbit Alexa555-conjugated secondary antibody (Thermo Fisher Scientific); Alexa405-coupled phalloidin (Invitrogen); Horseradish peroxidase-coupled anti-mouse and anti-rabbit IgGs (Invitrogen) were used for immunoblotting; Recombinant Human EGF Protein (R&D Systems). Erlotinib hydrochloride (500nM, Sigma-Aldrich); Dasatinib (500nM, Selleckchem); NSC-23766 (200μM, Tocris); CK-666 (100μM, Sigma-Aldrich); Latrunculin A (250nM, Sigma-Aldrich); SMIFH2 (20μM, Sigma-Aldrich); Jasplakinolide (100nM, Sigma-Aldrich); Y-27632 dihydrochloride (2-40μM, Sigma-Aldrich); Nocodazole (2-40μM, Sigma-Aldrich).

Four smartpool siRNA against EGFR was purchased from Dharmacon (Ref #: SO-2728829G). Targeted sequences were siRNA1, GCAUAGGCAUUGGUGAAUU; siRNA2, GCCAGGUCUUCAAGGAUGU; siRNA3, CCUUUAUGCUCCUCGGAAG; siRNA4, GAUUGGUGCUGUGCGAUUC. Four non-targeting pool siRNA (Dharmacon Ref#: SO-2736448G) was used as a nontarget control.

### Cell culture and transfection

S180-E-cad-GFP cells (Chu et al., 2004), a murine sarcoma cell line, were kindly provided by J. P.T. and tested for mycoplasma. S180-E-cad-GFP cells were cultured at 37°C, 5% CO2 in high-glucose Dulbecco’s Modified Eagle Medium (DMEM), supplemented with 10% fetal bovine serum (FBS) and 100IU/ml Penicillin/Streptomycin. For serum starvation, the cells were serum-starved in growth medium lacking FBS for more than 16 hours. Cells at 80% confluence were transiently transfected with 3 μg DNA using Neon electroporation system (Invitrogen). EGFR-mEOS and Actin-mapple were used for Neon transfection.

### Microwell preparation

The microwell was fabricated on both 12mm and 27mm glass bottom Iwaki dishes and visualized under the high-resolution spinning-disc confocal microscope. Master moulds with an array of 20-to 30-μm-diameter holes accompanied with 7 × 7 position markers were manufactured in SU8-3050 resist on silicon wafers using standard lithography techniques. The thickness of the SU8 resist layer sets the height of the microwells; here the total height was 46 μm. The details of the microwell preparation steps were described in (Engl et al., 2014).

### Imaging suspended cell-cell junction formation and analysis

After serum starvation, cells at 80% confluence were detached by flushing medium over the Petri dish to obtain a single-cell suspension. Cell seeding inside the microwell was achieved by adding one drop of prepared single-cell suspension. A glass-bottom dish (Iwaki) with suspended single cells inside microwells was mounted onto the microscope stage and immersed in serum-starved medium without Pheno red. Another drop of the single-cell suspension was added into the microwells for the second seeding (Supplementary Figure. 1a). Floating cells were removed after seeding. Images were acquired every 1 min using spinning-disc confocal microscopy.

As doublets are reorienting during spinning disc confocal microscopic imaging constantly, E-cad junction always presents at a certain inclination and is seldom in the focal plane. Therefore, a volume with 5-10 μm thick was imaged to ensure a thorough recording of the contact (Supplementary Figure. 1b). The details of the image processing procedures were described previously (Engl et al., 2014). With this processing, deconvoluted image stacks could be interpolated individually, enabling directly to visualize contact of cell doublets.

By following the evolution of junction radius in the function of time, we determined junction formation time (τ) by fitting the following exponential to the experimental junction formation curves (Figure. 2b and Supplementary Figure. 1g):

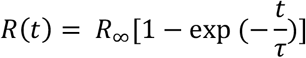

### Fluorescence recovery after photobleaching (FRAP)

The cortical actin recovery time (t_half_) was determined by fitting the following exponential to the recovery curves:

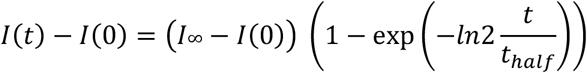

Two regions of interest were bleached for FRAP measurements: one on the cortex of singlets; another on the free cortex of doublets.

### Cortical tension measurement

Single S180-E-cad-GFP cells were blocked with 5% BSA and were deposited on a glass channel. Micropipettes were pulled with a flaming pipette-puller (P-2000, Sutter Instruments) and forged into 5-7 μm in diameter. The cortical tension here was computed with the law of Laplace: 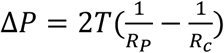, where *R*_*P*_ is the radius of the pipette, *R*_*c*_ is the radius of the cell and Δ*P* is the critical pressure when the extension of the surface of the cell into the pipette (*L*_*P*_) was equal to the radius of the pipette(*R*_*P*_). Thus, measuring cortical tension T is converted to measuring Δ*P*. Before each measurement, the pressure in the pipette was equilibrated with the outside pressure. When the pressure in the pipette was gradually decreased, cells will be aspirated into the pipette immediately. Once the cell formed a hemispherical protrusion in the pipette, which means *L*_*P*_ was equal to *R*_*P*_, the threshold pressure was reached.

### Western blotting

Confluent cells were lysed in a RIPA buffer (Sigma) with added protease and phosphatase inhibitor (Roche) for 20 min at 4 °C. Proteins extracted were quantified by BCA assay (Bio-Rad). 4-20% SDS polyacrylamide gel electrophoresis (PAGE) (Bio-rad) and eletro-transfer at 100V for 2h were performed to separate protein extracts and transfer them to PVDF membranes (Bio-Rad). Non-specific sites were blocked with 5% BSA in TBS 0.05% Tween 20. Membranes were incubated with primary antibodies (1:1000 dilution) overnight at 4 °C and then followed by an incubation of secondary HRP antibodies (1:2000) for 1h at room temperature. Results were visualized with ChemiDoc chemoluminescence detection system (Bio-rad). Quantification was performed using Image Lab. β-actin was used as a loading control to normalize the quantification.

### Immunofluorescent staining

After 1h junction formation, S180-E-cad-GFP cells were fixed with pre-warmed 4% paraformaldehyde in PBS at 37 °C for 15 min and then permeabilized with 0.2% Triton X-100 in TBS for 30 min at room temperature. Samples were blocked with 1% BSA in TBS for 1h, incubated with primary antibodies overnight at 4 °C and then incubated with secondary antibodies at room temperature for 1h.

### Data display and statistics

Prism (GraphPad Software) and Matlab (Math Works) were used for data analysis and graph plotting. Graphs were mounted using Adobe Illustrator. ANOVA test and paired or unpaired Student’s t-test were carried to analysis the significant difference levels.

## Notes

### Competing Interest Statement

The authors have declared no competing interest.

